# Genetic resistance to stripe rust infection of the wheat ear is controlled by genes controlling foliar resistance and flowering time

**DOI:** 10.1101/2021.04.27.441654

**Authors:** Laura Bouvet, Lawrence Percival-Alwyn, Simon Berry, Paul Fenwick, Sarah Holdgate, Ian J. Mackay, James Cockram

## Abstract

Yellow rust (YR), or stripe rust, is a fungal infection of wheat (*Triticum aestivum* L.) caused by the pathogen *Puccinia striiformis* Westend f. sp. *tritici* (*Pst*). While much research has focused on YR infection of wheat leaves, we are not aware of reports investigating the genetic control of YR resistance in other wheat structures, such as the ears. Here we use an eight-founder population to undertake genetic analysis of glume YR infection in wheat ears. Five quantitative trait loci (QTL) were identified, each explaining between ~3-7% of the phenotypic variation. Of these, three (*QYrg.niab-2D.2, QYrg.niab-4D.1* and *QYrg.niab-5A.1*) co-located with QTL for leaf YR resistance previously identified in the same population, with evidence suggesting *QYrg.niab-5A.1* may correspond to the adult plant resistance locus *Yr34* which originates from *T. monococcum* ssp. *monococcum* and that resistance at *QYrg.niab-2D.2* may be conferred by chromosomal introgression from a wheat relative. Additional leaf YR resistance QTL previously identified in the population were not detected as controlling glume resistance, with the remaining two glume YR QTL linked to genetic loci controlling flowering time. The first of these, *QYrg.niab-2D.1*, mapped to the major flowering time locus *Photoperiod-D1* (*Ppd-D1*), with the early-flowering allele from the MAGIC founder Soissons conferring reduced glume YR resistance. The second, *QYrg.niab-4A.1*, was identified in one trial only, and was located close to a flowering time QTL. This indicates earlier flowering results in increased glume YR susceptibility, likely due to exposure of tissues during environmental conditions more favourable for *Pst* infection. Collectively, our results provide first insights into the genetic control of YR resistance in glumes, controlled by subsets of QTL for leaf YR resistance and flowering time. This work provides specific genetic targets for the control of YR resistance in both the leaves and the glumes, and may be especially relevant in *Pst-prone* agricultural environments where earlier flowering is favoured.

**Core ideas:**

- *Puccinia striiformis* Westend f. sp. *tritici* (*Pst*) causes yellow rust (YR) in wheat leaves and ears.
- We present the first reports for the genetic control of YR on the wheat ear.
- Ear YR infection is controlled by subsets of QTL controlling leaf resistance and flowering time.
- The findings are relevant to wheat breeding for *Pst*-prone environments.

## Introduction

The fungal pathogen *Puccinia striiformis* Westend f. sp. *tritici* (*Pst*) causes yellow rust (YR), also known as stripe rust, in wheat (*Triticum aestivum* L.) (reviewed by Bouvet et al. 2021a). The visible symptoms of YR infection, such as sporulation, are typically observed on wheat leaves. However, under the right conditions *Pst* urediniospores (thin walled spores produced during the asexual stage of the *Pst* lifecycle) will also colonise leaf sheaths, as well as structures of the wheat ear including the outermost structures that surround each floret (the glumes) (Figure 1a-b). While the main concern for wheat growers is foliar infection, glume infection can become a significant problem during moderate to severe epidemics (Bouvet et al. 2021a). *Pst* urediniospores infect floret tissues from heading date (Zadoks Growth Stage 55, GS55; Zadoks et al. 1974) until flowering (GS61), with symptoms appearing 10 to 20 days later (Wellings, 2003). Inside each floret, *Pst* germination and sporulation will occur on glumes, lemma and palea. At a first glance, infected wheat heads appear discoloured or bleached, and symptoms can be mistaken for that of other diseases such as Fusarium head blight. Upon closer inspection, pustules inside the spikes are more evident and opening the floret exposes urediniospores, thereby confirming *Pst* infection. These symptoms are shortlived compared to foliar infection, mainly because cycles or re-infection are more difficult when spores are encased within the florets, and because they occur towards the end of the plant lifecycle, including fertilization and grain filling. An additional symptom includes shrivelled grains, although this is largely dependent on the timing on infection. Generally, the earlier the infection at the onset of flowering, the higher the extent of shrivelled grain. In severe cases, glume infection can lead to grain quality downgrading, as exemplified by reports of a 77% reduction in grain weight relative to uninfected plants (Cromey, 1989a) as well as a 20% yield reduction in a resistant variety (Purdy & Allan, 1965). *Pst* glume infection is mostly observed in varieties which are moderately to highly susceptible to YR infection of the leaf (Wellings, 2003). Thus, a variety with foliar resistance will generally also show resistance in the ear, while a susceptible variety is much more likely to exhibit symptoms of glume infection. However, there have been reports of moderately resistant varieties exhibiting unusually high levels of glume infection (Cromey, 1989b; Purdy & Allan, 1965; Wellings, 2003, 2009). While little is known about the effects of environmental conditions, ear morphology and the timing of flowering on YR glume infection, reports suggest that moisture and cool temperatures at flowering are key factors in the onset of significant outbreaks, and that florets are least susceptible after anthesis (Cromey, 1989a).

**Figure 1.**
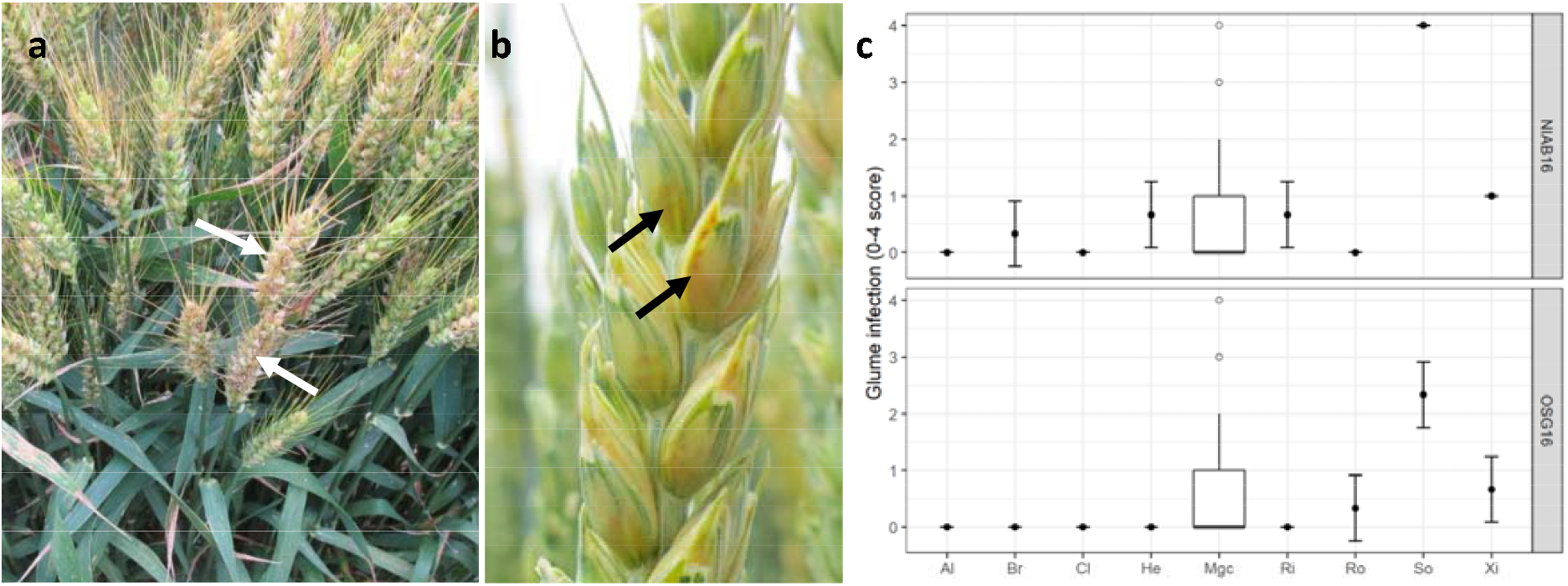
MAGIC founder yellow rust (YR) infection of the ear. Examples of founder Soissons at (A) the plot level, and (B) a close-up of a single ear. Arrows indicate example areas of notable infection, highlighting notable infections of the ear in general (white arrows) and the glumes (black arrows). (C) Distribution of glume YR infection of the MAGIC founders, and the MAGIC recombinant inbred lines (RILs) at the trials NIAB16 and OSG16. Mean glume infection is represented by filled black dots for each fonder, with standard deviation plotted as error bars. The central boxplot represents the distribution of the MAGIC RILs (single counts for un-replicated lines and total counts for replicated lines). Black unfilled circles represent outliers. MAGIC founders are listed as, Al: Alchemy, Br: Brompton, Cl: Claire, He: Hereward, Mgc: MAGIC RILs, Ri: Rialto, Ro: Robigus, So: Soissons, Xi: Xi19.

*Pst* infection of the leaves prevents leaf tissue from photosynthesizing. Extensive infection of the ears may have a similar detrimental effect on photosynthesis, particularly when leaf photosynthetic area is already significantly reduced due to extensive sporulation, necrosis and/or chlorosis. Ear photosynthesis significantly contributes towards grain filling by providing the developing grain with an important source of photoassimilates (10-76 %) and has been shown to exhibit similar levels of CO_2_ fixation to the flag leaf (Maydup et al. 2010; Tambussi et al. 2007). These characteristics are particularly relevant under drought conditions, when ears exhibit higher water use efficiency than flag leaves (Tambussi et al. 2007). The photosynthetic capacity of ears has therefore been recognised as an area relevant to wheat breeding and is the target of further optimisation as part of international research efforts looking to increase wheat yield potential, such as the Wheat Yield Consortium (Parry et al. 2011). In addition, if we consider the ways in which *Pst* urediniospores can colonise plant tissue, glume infection could potentially pose a threat to hybrid wheat breeding: as wheat is a predominantly inbreeding crop, developing hybrids requires male sterility in female parents, which receive pollen from male parents. This critical cross-pollination step relies on open florets during anthesis, providing an easy entry point for *Pst* urediniospores. In the context of a moderate to severe yellow rust outbreak, glume infection could result in increased levels of shrivelled grain, posing a serious problem for the large-scale seed multiplication efforts required by commercial hybrid breeding.

More than 300 genetic loci controlling resistance to foliar YR infection have been identified, of which around 80 represent permanently named yellow rust resistance (*Yr*) genes (summarised by Jamil et al. 2020). However, the genetic components underlying glume resistance have not been explored, with current knowledge essentially limited to its phenotypic symptoms. Recently, several multi-founder experimental populations have been developed in wheat (reviewed by Scott et al. 2020). Here we use an eight-parent wheat multi-founder advanced generation inter-cross (MAGIC) population genotyped using a 90,000 feature single nucleotide polymorphism (SNP) array which has previously been used for genetic mapping of YR infection of the leaf (Bouvet et al. 2021b) to: (1) characterise for the first time the genetic basis of glume YR resistance, and (2) compare glume resistance QTL to the genetic basis of yellow rust resistance in leaves.

## Methods

### Wheat germplasm and field trials

The eight-founder ‘NIAB Elite MAGIC’ wheat population is previously reported (Mackay et al. 2014). Briefly, it consists of eight founders (the varieties Alchemy, Brompton, Claire, Hereward, Rialto, Robigus, Soissons and Xi19) inter-crossed over three generations, after which >1,000 recombinant inbred lines (RILs) were developed by inbreeding. A subset of the population (n=552) was sown in two untreated, partially replicated trials sown in the autumn of 2015, and grown to maturity to the following summer of 2016. The trial sites were in the United Kingdom (UK) at NIAB in Cambridgeshire (NIAB16; latitude 52.235010, longitude 0.097871) and Osgodby in Lincolnshire (OSG16; latitude 53.410161, longitude -0.386770). Each trial consisted of 1,200 plots arranged in 20 columns and 60 rows. Trial design consisted of 2 main-blocks, with 3 blocks and 2 sub-blocks per block, containing 444 MAGIC RILs in two replicates and 234 un-replicated RILs. Trial designs were generated using DEW experimental design software (www.expdesigns.co.uk). Each RIL was sown in two 1-m rows, with six rows sown adjacent to each other at a time, thus forming 1×1m areas consisting of three RILs and/or founders. At NIAB16, the central two rows were sown with the YR susceptible variety, Vuka, which served to help spread YR infection. In contrast, at OSG16, 1×1 m plots of Vuka were sown every three traverses. NIAB16 was artificially inoculated with a mixture of *Pst* races: ‘Solstice’ (isolate 08/21 virulent on *Yr 1, 2, 3, 4, 6, 9, 17, 25, 32*) and ‘Warrior’ (isolate 11/08 virulent on *Yr 1, 2, 3, 4, 6, 7, 9, 17, 25, 32, Sp*). For OSG16, no inoculation was undertaken. Both trials were subject to very high levels of natural *Pst* inoculum early in the season, and before artificial inoculation had taken place, and it is therefore highly likely that both sites were predominantly exposed to the effects of natural *Pst* infection. Based on UK Cereal Pathogen Virulence Survey (UKCPVS) data of *Pst* isolates in 2016, Cambridgeshire and Lincolnshire (including samples from the Osgodby site) were dominated by the Warrior group of races (UKCPVS, 2017) (Supplementary Table 1).

### Glume YR assessment and phenotypic data analysis

Glume infection was assessed at the late milk development stage (GS77). For each plot, 10-12 ears were selected at random and the number of spikelets per ear exhibiting glume infection counted. This was converted into a percentage to obtain an estimate of the proportion of ear infection, which was assigned a number on a five-point scale from 0-4, where 0 represents no glume infection, 1 = 25% of glumes infected, 2 = 50% of glumes infected, 3 = 75% of glumes infected, and 4 represents all glumes per head infected (examples shown in Supplementary Figure 1). A score was allocated when ≥50 % of the ears within a plot corresponded to that score. Linear mixed models based on Restricted Maximum Likelihood (REML) were used to adjust the infection data and take in account any spatial variation. Best Linear Unbiased Estimators (BLUEs) were computed following a step-wise model selection approach. Baseline models considered were as follows: Model 1 – Blocking: genetic effects are estimated based only on the inter- and intra-block variation recovered from the model. Model 2 – spatial: only considers global and/or local field trends. Model 3 - spatial + blocking: combination of the above models. Models were fitted using Genstat 18^th^ edition (VSN International, 2015). Akaike Information Coefficient (AIC) was used as a measure for model selection (Akaike, 1974): each model was optimised using AIC as a measure of model fit, and the model with the lowest AIC value selected. Inspection of the glume infection distribution and residual plots revealed a dataset skewed towards resistance. Therefore, the natural log was applied to improve the normality of the data. Phenotypic data for YR infection of the leaf in the NIAB16 and OSG16 trials has previously been reported, as part of a wider series of five trials investigating the genetics of foliar YR infection (Bouvet et al. 2021b). To determine the consistency of the phenotype across test environments, and the relationship between YR glume infection with YR leaf infection previously reported in the same trials (Bouvet et al. 2021b), Pearson correlation coefficients for glume infection were computed using the ‘cor.test()’ function in the statistical software R 3.6.1 (R Core Team, 2019).

### Genetic analyses and bioinformatics

The MAGIC population was previously genotyped using a 90,000 feature Illumina SNP array (Wang et al. 2014), as described by Mackay et al. (2014), and a MAGIC genetic map subsequently generated (Gardner et al. 2016). QTL analysis was undertaken following four different statistical methods. Method 1 - single marker analysis (SMA): regression analysis on allelic state of 7,369 mapped SNP markers from the MAGIC genetic map. The remaining analyses were based on founder haplotype probabilities calculated with the ‘mpprob’ function in R/mpMap (Huang et al. 2011) with a threshold of 0.5. Method 2 – identity by descent (IBD): regression analysis on founder haplotype probabilities using R/qtl. Method 3 - interval mapping (IM): conducted in R/mpMap using the haplotype probability estimates. Method 4 - composite interval mapping (CIM): as IM but with 10 marker covariates. For IM and CIM, the empirical *p* = 0.05 significance threshold was computed in R/mpMap using the ‘sim.sig.thr’ function, producing a cut-off at *p* = 1.6E-05 (−log10(*p*) = 4.8) for NIAB16 and *p* = 3.7E-05 (−log10(*p*) = 4.2) for OSG16. For SMA and IBD, the package R/qvalue was used to correct for multiple testing, with a *q*-value threshold of 0.05 (R Core Team, 2017). QTL intervals were defined as a genomic region which, at a given genetic position had a *q-*value ≤1.6E-05 (NIAB16) and *q* ≤ 3.7E-05 (OSG16). The genetic versus physical map location of each QTL was determined for each relevant chromosome by plotting the location of SNPs on the MAGIC genetic map (Gardner et al. 2016) against their position on the physical map of the wheat reference genome assembly (RefSeq v1.0; IWGSC et al. 2018) and the location of the most significant SNP for a given QTL highlighted. Data were plotted using R/ggplot2 (Wickham, 2009).

To investigate the genomic regions that span the QTL found to confer yellow rust resistance in both the glumes and leaf (Bouvet et al. 2021b), we undertook three analyses. (1) Sequence alignment depth analysis: previously generated Illumina paired-end sequence reads from wheat cultivars Cadenza, Claire, Chinese Spring, Paragon, Robigus and Weebill1 (Clavijo et al. 2017; Walkowiak et al. 2020) were trimmed using trim-galore (Krueger 2019) to remove adapter sequences and quality filter (-q 25), retaining only reads with a length greater or equal to 100 bp before aligning using BWA-MEM (Li 2013) against the cv. Chinese Spring wheat reference genome assembly (RefSeq v1.0, IWGSC 2018). Following BWA-MEM alignment, SAMtools fixmate was first used to fix and fill in mate information before filtering to retain proper read pairs and remove non-primary alignments using SAMtools view (Li et al. 2009). The alignments were then sorted with SAMtools before removing sequencing duplicates using Picard MarkDuplicates (http://broadinstitute.github.io/picard). BAM files for each variety were then merged and indexed (-c) using SAMtools. A bespoke python script ‘bam_depth.py’ which uses pysam (https://github.com/pysam-developers/pysam) was used to extract the mean average sequencing depth and coverage for 5 kb regions every 2 kb along the three target chromosomes (2D, 4D and 5A). Another bespoke python script ‘normailse.py’ was used to calculate median average depth from 5 kb regions with 100% sequence coverage for each variety before normalising the depth of all 5 kb regions to this average. Finally, a sliding (every 0.05 Mbp) window (0.5 Mbp in length) was used to extract the median average from all the 5 kb regions within the window, and results plotted using matplotlib (Hunter, 2007). (2) Sequence divergence analysis: to investigate the cause of large regions of very low BWA-MEM alignment depth identified in step-1, we used the same MiniMap2 (Li, 2018) alignment parameters that Liftoff (Shumate & Salzberg, 2020) uses for annotating closely related species. Gene-size regions (3 kb, subsequently termed ‘genes’) every 20 kb along the cv. Chinese Spring wheat reference genome assembly (RefSeq v1.0; IWGSC, 2018) were extracted using samtools faidx (Li et al. 2009) and mapped to the assemblies of cvs. Cadenza, Chinese Spring, Claire, Robigus, Paragon and Weebill1 (Clavijo et al. 2017, Walkowiak et al. 2020) using the parameters -a --end-bonus 5 --eqx -N 50 -p 0.5. The mean average gap-compressed per-base sequence divergence for primary alignments were calculated for sliding (every 2 Mbp) windows (50 Mbp in length) using the original chromosome level assembly for the sequential positions of the gene-size regions. (3) Depth analysis of unmapped reads: to rule out deletions causing regions of extremely low mapping depth in step-1 above and the associated sequence divergence in step-2 being caused by mapping to homoeologous genes/regions, unmapped reads from step-1 were investigated. Unmapped reads were extracted from the Chinese Spring sequence alignment files using samtools fastq. Single reads and read pairs were mapped back to their own assemblies. Using the alignment files generated in the sequence divergence analysis, unique scaffolds were identified within a 1 Mbp region of Chinese Spring, and the self-alignment depths of these scaffolds were averaged and plotted every 0.5 Mbp.

## Results

### Pst glume infection

Genotyped MAGIC lines were assessed for YR disease severity in the ears (Figure 1a-b) at two test environments in 2016: trials NIAB16 and OSG16. Amongst the eight MAGIC founders, cv. Soissons showed high glume YR infection in both trials (mean >2.5; Figure 1c). The remaining seven founders were found to be relatively resistant, with mean glume infection (GIF) scores ≤1, with Xi19 showing the second highest founder glume infection in both trials. The MAGIC RILs showed phenotypic variation over the full range of the assessment scale (Figure 1c): nearly 70 % of RILs showed little or no infection (GIF = 0) while ~10 % exhibited *Pst* infection in the majority of the ear (GIF = 3-4) (Supplementary Figure 2). Pearson’s correlation coefficient analysis of the MAGIC glume YR infection data alongside previously reported leaf YR infection data from the same trials (Bouvet et al. 2021b) found a positive correlation at both environments, with OSG16 (*R* = 0.62, *p* <2.2E-16) showing a stronger correlation than was observed at NIAB16 (*R* = 0.49, *p* < 2.2E-16) (Figure 2). MAGIC RILs susceptible in the glumes generally also exhibited high leaf susceptibility (leaf infection >60%). However, a few notable exceptions were observed, particularly at NIAB16 where six RILs (accessions 5954, 6216, 5770, 5883, 6110, 6374) were found to be moderately resistant on leaves (infection score ≤25%) but susceptible in the glumes (GIF >3.5). One of these, RIL 5954 displayed the same phenotypic combination at OSG16. All glume YR phenotypic data is listed in Supplementary Table 2.

**Figure 2.**
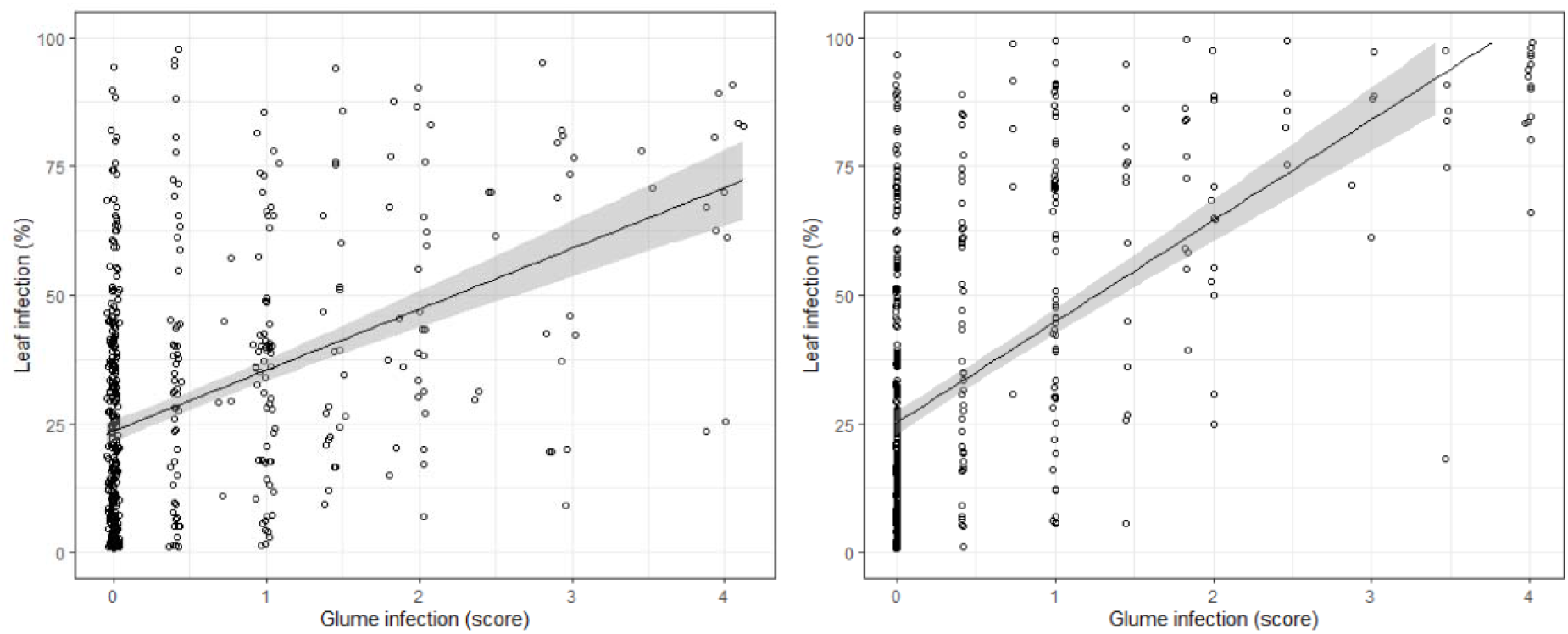
Correlation between MAGIC recombinant inbred line (RIL) yellow rust infection of the glume and the leaf at trials NIAB16 (left) and OSG16 (right). Back-transformed adjusted means were used for both scatter plots. The black line is the linear regression line and the grey area the 95 % confidence level interval. Pearson correlation coefficients: NIAB16: *R* =0.49, *p* <2.2E-16; OSG16: *R* = 0.62, *p* <2.2E-16. Yellow rust leaf phenotypic data was sourced from Bouvet et al. (2021).

Statistical analysis of the glume infection phenotypic data using REML showed that the Wald statistic for genotype effect was highly significant (Chi pr <0.001) at both environments (Supplementary Table 3). High genotypic performance is further supported by high broad sense heritability values at NIAB16 (*h^2^* = 0.80) and OSG16 (*h*^2^ = 0.82), mirroring values observed for leaf infection at the same trials (*h*^2^ = 0.79-0.94) (Bouvet et al. 2021b). This evidence points towards stable genotypic performance for glume infection in the MAGIC population at each site. However, the weaker correlation of glume infection scores between OSG16 and NIAB16 (*R* = 0.47, *p* <2.2E-16) highlights possible genetic x environmental interaction (Supplementary Figure 3), or could be due to difference in the *Pst* pathotypes present at each site.

### QTL mapping of glume YR resistance and comparison with leaf resistance QTL

Five glume YR resistance QTL were identified, located on chromosomes 2D (*QYrg.niab-2D.1, QYrg.niab-2D.2*), 4A (*QYrg.niab-4A.1*), 4D (*QYrg.niab-4D.1*) and 5A (*QYrg.niab-5A.1*), as listed in Table 1, with MAGIC founder effects at each QTL summarised in Table 2. The percentage of the phenotypic variance explained (PVE) for each QTL was around 5%, ranging from 3.4 to 6.8%, and jointly accounting for ~23% PVE. For each QTL, placing the peak SNP on plots of genetic versus physical map positions for all other markers on the corresponding chromosomes allowed QTL locations to be viewed in genomic context (Figure 3). Four of the five QTL were located on the long arms of their respective chromosomes in regions of high genetic recombination, ≤43 Mbp from the telomere. Indeed, *QYrg.niab-4D.1* (4.8% PVE), *QYrg.niab-2D.2* (3.9% PVE) and *QYrg.niab-5A.1* (6.8% PVE) were located very close to the ends of their chromosomes, just 7, 13 and 19 Mbp from the chromosome 4D, 2D and 5A telomeres, respectively. *QYrg.niab-4A.1* (3.4% PVE) was ~39 Mbp from the chromosome 4A long arm telomere and was located within a region of low agreement between SNP genetic versus physical map order, and spanning a stretch of chromosome 4A from 170-210 cM previously identified as a putative introgression from the related wheat species *Triticum dicoccoides* present in the MAGIC founder Robigus (Gardner et al. 2016). However, our results indicate the glume YR resistance allele at *QYrg.niab-4A.1* is conferred by Xi19, and so does not originate from the chromosome 4A alien introgression present in Robigus. *QYrg.niab-2D.1* (6.4% PVE) was located halfway down the chromosome 2D short arm in a region of high genetic recombination.

**Figure 3.**
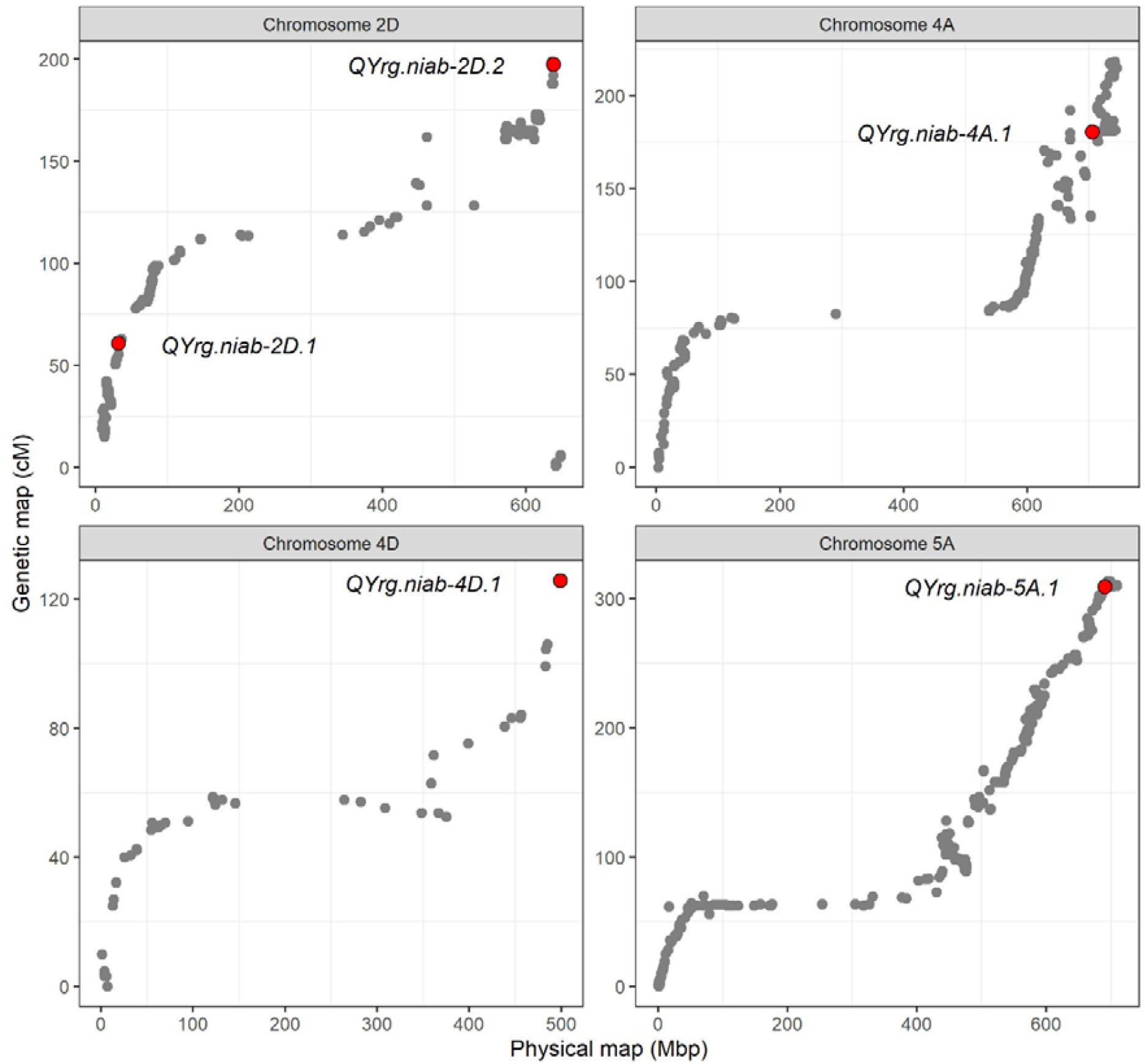
Genomic locations of the five quantitative trait loci (QTL) conferring glume yellow rust resistance in the MAGIC population. For each chromosome, plotted are the physical (Mbp; IWGSC, 2018) versus genetic (cM; Gardner et al. 2016) map positions of the genetic markers, with the peak marker locations for each QTL shown in red.

**Table 1.**
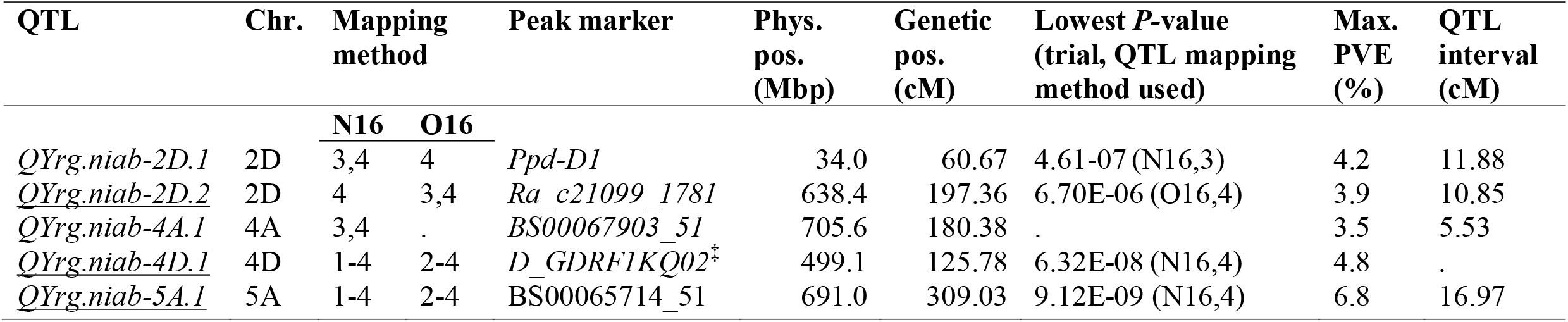
Glume yellow rust (YR) resistance quantitative trait loci (QTL) identified in trials NIAB16 (N16) and OSG16 (O16). Glume QTL colocating with previously identified leaf QTL (Bouvet et al. 2021b) are underlined. QTL mapping methods are numbered as: (1) single marker analysis, (2) identity by descent, (3), interval mapping, and (4) composite interval mapping with 10 covariates. The lowest *p*-values identified for each QTL, sourced either from trial N16 or O16, are indicated. In a similar way, values for the maximum percentage of the phenotypic variance explained (% PVE) by each QTL are indicated. Physical and genetic map information is from IWGSC (2018) and Gardner et al. (2016), respectively. QTL detected by method-4 only across both environments were not included here. ^‡^Full marker name: *D_GDRF1KQ02H66WD_341*.

| QTL | Chr. | Mapping method |  | Peak marker | Phys. pos. (Mbp) | Genetic pos. (cM) | Lowest *P*-value (trial, QTL mapping method used) | Max. PVE (%) | QTL interval (cM) |
| --- | --- | --- | --- | --- | --- | --- | --- | --- | --- |
|  |  | N16 | O16 |  |  |  |  |  |  |
| <u>*QYrg.niab-2D.1*</u> | 2D | 3,4 | 4 | *Ppd-D1* | 34.0 | 60.67 | 4.61-07 (N16,3) | 4.2 | 11.88 |
| <u>*QYrg.niab-2D.2*</u> | 2D | 4 | 3,4 | *Ra_c21099_1781* | 638.4 | 197.36 | 6.70E-06 (O16,4) | 3.9 | 10.85 |
| <u>*QYrg.niab-4A.1*</u> | 4A | 3,4 | . | *BS00067903_51* | 705.6 | 180.38 | . | 3.5 | 5.53 |
| <u>*QYrg.niab-4D.1*</u> | 4D | 1-4 | 2-4 | *D_GDRF1KQ02*<sup>‡</sup> | 499.1 | 125.78 | 6.32E-08 (N16,4) | 4.8 | . |
| <u>*QYrg.niab-5A.1*</u> | 5A | 1-4 | 2-4 | *BS00065714_51* | 691.0 | 309.03 | 9.12E-09 (N16,4) | 6.8 | 16.97 |

**Table 2.** Predicted founder effects at the five quantitative trait loci (QTL) conferring glume yellow rust resistance, based on natural-log adjusted BLUEs. For each QTL, founder effects presented are for the consensus marker listed in Table 1 with the most significant *p*-value identified with QTL mapping method-3 (interval mapping) and method-4 (composite interval mapping with 10 covariates). MAGIC founders, Al: Alchemy, Br: Brompton; Cl: Claire, He: Hereward, Ri: Rialto, Ro: Robigus, So: Soissons, Xi: Xi19. Founder effects are all relative to those of Xi19.

| QTL | Chr. | Predicted founder effects |  |  |  |  |  |  |  | Trial, QTL method |
| --- | --- | --- | --- | --- | --- | --- | --- | --- | --- | --- |
|  |  | Al | Br | Cl | He | Ri | Ro | So | Xi |  |
| *QYrg.niab-2D.1* | 2D | 0.09 | 0.01 | -0.05 | -0.04 | 0 | 0 | 0.3 | 0 | NIAB16, 3 |
| *QYrg.niab-2D.2* | 2D | -0.24 | 0.09 | -0.05 | 0.43 | -0.32 | -0.26 | 0.32 | 0 | NIAB16, 4 |
| *QYrg.niab-4A.1* | 4A | -0.27 | -0.18 | -0.2 | -0.34 | -0.33 | -0.32 | -0.41 | 0 | NIAB16, 3 |
| *QYrg.niab-4D.1* | 4D | 0.23 | -0.08 | NA | 0.32 | 0.55 | 0.52 | 0.4 | 0 | OSG16, 3 |
| *QYrg.niab-5A.1* | 5A | -0.18 | -0.12 | -0.15 | -0.31 | 0.07 | -0.1 | 0.13 | 0 | OSG16,4 |

Our previous study reported leaf YR resistance QTL at five sites using the MAGIC population, which included the NIAB16 and OSG16 trials investigated here for glume YR resistance (Bouvet et al. 2021b). Cross-comparison between the glume and YR resistance QTL showed three of our glume QTL to co-locate with leaf QTL: (1) glume QTL *QYrg.niab-2D.2* and leaf QTL *QYr.niab-2D.1* (~18% PVE; Bouvet et al. 2021b), both identified by the same peak marker (*Ra_c21099_1781*). (2) Glume QTL *QYrg.niab-4D.1* and leaf QTL *QYr.niab-4D.1* (4% PVE; Bouvet et al. 2021b), both defined by the same peak marker (*D_GDRF1KQ02H66WD_341*). (3) Glume QTL *QYrg.niab-5A.1* and leaf QTL *QYr.niab-5A.1* (3.6% PVE; Bouvet et al. 2021b), for which the peak markers were 8 cM apart. For the remaining two glume resistance QTL, the peak marker for *QYrg.niab-2D.1* was a diagnostic marker within the *Photoperiod-D1 (Ppd-D1*) gene, able to distinguish the early flowering photoperiod insensitive *Ppd-D1a* allele carried by Soissons from the photoperiod sensitive *Ppd-D1b* alleles carried by the remaining seven founders (Bentley et al. 2013). The predicted allelic effects for QTL *QYrg.niab-2D.1* showed increased glume YR susceptibility was conferred by the Soissons allele at this locus. Finally, *QYrg.niab-4A.1*, identified in NIAB16 only and explaining 3.4% PVE, was not found to co-locate with any QTL for yellow rust resistance in leaves. Glume QTL *QYrg.niab-4A.1* did not co-locate with any of the MAGIC leaf YR resistance QTL identified by Bouvet et al. (2021b).

Finally, we cross-referenced our three glume/leaf resistance QTL with previously named *Yr* resistance loci. *QYrg.niab-2D.2/QYr.niab-2D.1* overlapped with named adult plant resistance gene *Yr54* (Banset et al. 2014), which is located downstream of the genetic marker gwm301 (at 642.3 Mbp on chromosome 2D in the wheat reference genome). For *QYrg.niab-4D.1/QYr.niab-4D.1* (peak QTL position at 499.1 Mbp in the reference wheat genome), three named *Yr* genes are reported on chromosome 2D: *Yr22* (Chen et al. 1995), *Yr46* (Moore et al. 2015) and the provisionally designated *YrS2* (Sun et al. 2019). *Yr22* has not been mapped intrachromosomally, while the adult plant resistance gene *Yr46* is located ~80 Mbp away. However, based on flanking marker cfd84 (4D: 498.7 Mbp), *YrS2* is within our glume/leaf rust resistance QTL interval. Finally, the end of the long arm of bread wheat chromosome 5A (between ~694-709 Mbp in the reference wheat genome) is recently reported to contain an introgression of chromosome 5A^m^ from the diploid wheat relative *T. monococcum* ssp. *monococcum* (Chen et al. 2021). This region of chromosome 5A^m^ carries the adult plant resistance locus *Yr34*, and its physical map location in the reference wheat genome places it within the physical map interval of our glume/leaf QTL *QYrg.niab-5A.1/QYr.niab-5A.1*. To begin to investigate the possible relevance of alien introgression to our glume/rust YR QTL, we undertook three bioinformatic analyses using publicly available data. First, we aligned Illumina paired-end genomic sequence data for six wheat varieties (which included the MAGIC founders Claire and Robigus) to the wheat cv. Chinese Spring reference genome. This identified regions of chromosomes 2D and 5A for which sequence reads from the six investigated wheat varieties did not align to the wheat reference genome when using the standard BWM mem alignment thresholds, while for chromosome 4D no such large-scale regions of non-alignment were identified (Figure 4). Mid-to-large scale regions of nonalignment on chromosomes 2D (~635.5-641.0 Mbp based on the positions in the reference genome) and 5A (~703.0-708.0 Mbp) were found to coincide with the QTL intervals of our glume/rust resistance loci *QYrg.niab-2D.2/QYr.niab-2D.1* and *QYrg.niab-5A.1/QYr.niab-5A.1*, respectively. To investigate whether these regions of non-alignment represented chromosomal deletions or putative alien chromosomal introgressions, we used the same alignment thresholds used for the liftover annotation of related species, to map uniformly distributed (every 20 kb) gene size (3 kb) regions of the Chinese Spring reference to the assemblies of the six wheat varieties. We then calculated the average sequence diversity of ‘genes’ from the six wheat varieties relative to their equivalent genes in the reference assembly (Figure 4). This showed that the ‘gene’ space within the regions of non-alignment identified in step-1 that overlapped with the locations of our YR resistance QTL on chromosomes 2D and 5A to both contain higher than expected levels of sequence variation relative to the reference genome, thus indicating possible alien introgression at both loci. No signatures of alien chromosomal introgression were identified for our chromosome 4D glume/leaf YR resistance locus. Finally, to provide further evidence the ‘gene’ diversity identified in our QTL regions on 2D and 5A in step-2 were not due to alignment of homoeologous/paralogous sequences, we took all the unmapped reads from step-1 and mapped these back to their own assemblies. Self-alignment depths were averaged and anchored back to the Chinese spring reference using the alignment files from our diversity analysis. The resulting increased read depth in the QTL regions on 2D and 5A indicated that lack of coverage in step-1 was due to the presence of sequences with lower sequence homology to the reference genome (characteristic of alien introgression), rather than chromosomal deletions (Supplementary Figure 4).

**Figure 4.**
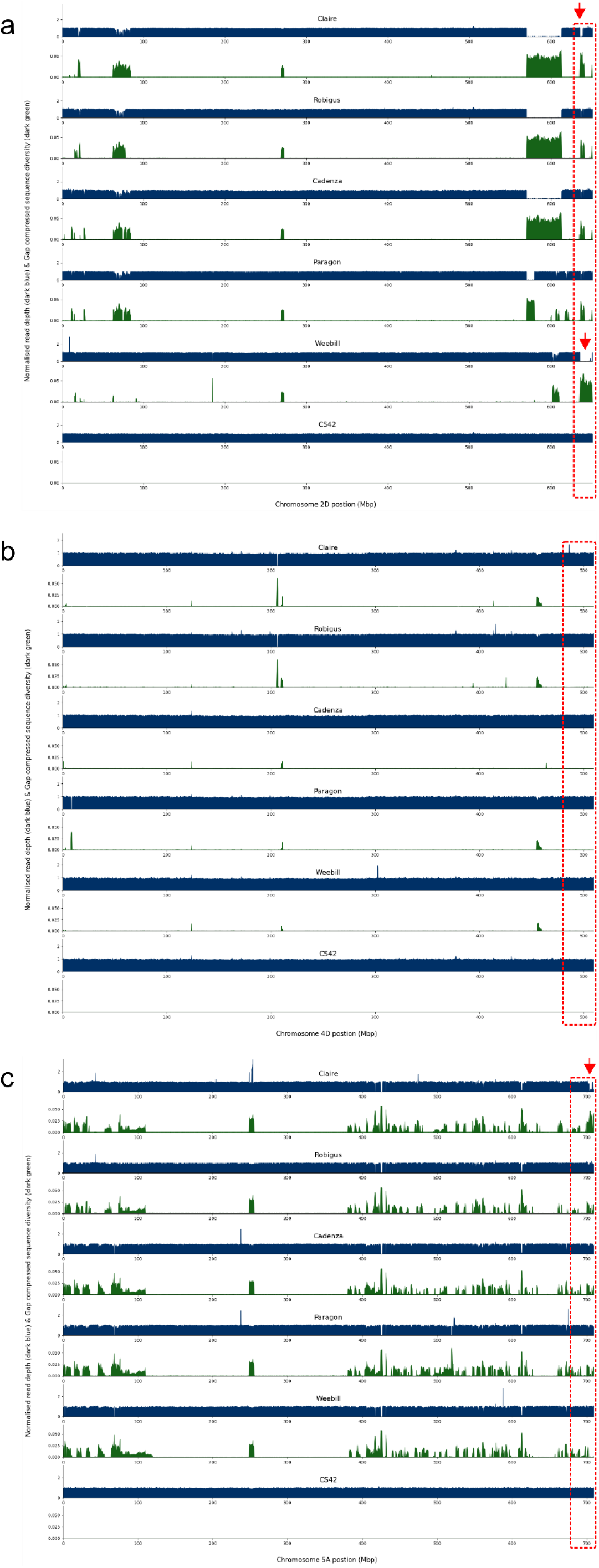
Signatures of alien chromosomal introgression in the genomes of six wheat cultivars: the MAGIC founders Claire and Robigus, as well as cvs. Cadenza, Paragon, Weebill1 and Chinese Spring (CS). Data for chromosomes (a) 2D, (b) 4D, and (c) 5A are shown. In each of these three panels, two data tracks are displayed per cultivar. The top track (dark blue) shows DNA sequence read coverage depth for each specific cultivar, as aligned to the CS reference genome using BWA-MEM, an alignment tool commonly used to map wheat sequences to the correct chromosome. The lower (dark green) track shows DNA sequence diversity of ‘genes’ (more accurately, gene-sized regions of 3 kb) derived from the CS reference genome and aligned against each specific variety using thresholds suitable to map sequences from closely related cereal species. The yellow rust resistance QTL intervals are indicated by the red dashed box. Putative regions of alien chromosomal introgression within these QTL intervals are indicated with a red arrow.

## Discussion

Wheat ear photosynthetic capacity plays a significant role in grain filling (Wang et al. 2016), particularly under stress conditions (Tambussi et al. 2007). While ear YR infection may not currently be as much of a recurrent problem as leaf infection, at least in the European context, it can become detrimental to yields provided suitable growing and environmental conditions. Due to climate change, temperatures are predicted to rise in many of the wheat growing areas of the world (IPCC, 2019). Such changes, including during the important wheat developmental stages of ear emergence, flowering and early grain fill, may result in wheat pests and diseases posing a greater, or changing, threat to optimum wheat grain yields. Indeed, simulations of UK wheat development under greenhouse gas emission scenarios finds acceleration of days to anthesis to accelerate by 13-19 days (Harkness et al. 2020). Despite such current and future risks, no previous reports of the genetic basis of wheat ear YR resistance have been published. Here, we begin to address this knowledge gap, finding that glume YR infection is controlled by a subset of genetic loci that confer resistance to foliar YR infection, combined with pleiotrophic effects of loci controlling the timing of ear emergence and subsequent development. Below we explore in more detail some of the issues related to these findings.

### Glume YR infection and phenotype

Variation for ear infection in the MAGIC population was skewed towards resistance, with the majority of MAGIC RILs showing little-to-no yellow rust infection. This trend is different to that observed for foliar infection reported in the same trials, which exhibited a less skewed range of phenotypic variation (Bouvet et al. 2021b). Assessing the extent of yellow rust infection in ears is harder compared to foliar infection, as disease symptoms are more easily and rapidly identified on leaves. Here, the degree of difficulty in phenotypic assessment had to be balanced with the practicalities of scoring large numbers of plots in the field. Therefore, a simple 0-4 scale was developed, which would have inevitably grouped together finer levels of response. Notably, infection may have been missed completely if it was confined to the inner layers of the floret which are not visible without manual intervention. This may partly explain the relatively low correlation between leaf and glume infection (NIAB16 *R* = 0.49; OSG16 *R* = 0.62), and on the power and precision to identify QTL. Leaf and ear infection not only differed in phenotypic distribution, but also in broad sense heritability: ear infection exhibiting >10% lower heritability in both environments, indicating additional non-genetic factors have a larger impact on the phenotypic response in ears than in leaves, and/or that the limitations of the 0-4 glume infection scale combined with the inherent difficulty in accurately scoring glume infection impacted phenotyping accuracy. This difference in heritability is in agreement with YR resistance studies in Triticale (a hybrid between wheat and rye) that also report lower heritability in ears compared to leaves (Losert et al. 2017). Furthermore, Losert et al. (2017) argue that the severe ear infection they observed in Triticale may have been a feature of the prevalent *Pst* ‘Warrior’ race and its potential ability to develop at warmer temperatures. Although temperature adaptation of the ‘Warrior’ race variants has not been experimentally confirmed, as they have for the aggressive race *PstS2* for example (Milus et al. 2009), it is plausible that warmer temperatures not previously conducive of germination and subsequent colonisation enable the pathogen to prolong its growth into photosynthetic tissue later in the season, such as the glumes (Mboup et al. 2012). The UK regions in which the trials were conducted were dominated by the ‘Warrior’ group of *Pst* races (Hubbard et al. 2017; Supplementary Table 1). Indeed, 2016 was a year that saw a high number of yellow rust infected leaf samples received by the UK Cereal Pathogen Virulence Survey, with some samples received as early as January (Hubbard et al. 2017). In addition to inoculum levels, other factors may explain between site differences in glume YR infection. For example, climatic differences may have played a role, with the OSG16 trial located in a cooler and wetter region than the NIAB16 trial. This likely led to a longer window of infection for *Pst* urediniospores, making disease symptoms more pronounced and easier to score later in the season, thus resulting in a stronger correlation between leaf and ear infection. Overall, our observations were in line with reports that ear YR infection is prominent when significant levels of inoculum are present and are coupled with suitable growing conditions and timely flowering (Cromey, 1989a; Wellings, 2003).

### The role of foliar YR resistance loci in glume resistance

The co-location of three of our five glume YR QTL with foliar resistance QTL previously identified in the same trials (Bouvet et al. 2021b) indicates the presence genetic mechanisms in common. Based on the percentage of the phenotypic variance explained, the leaf YR QTL identified in the MAGIC population by Bouvet et al. (2021b) were classified as either ‘major’ (>5% PVE, 8 QTL) or ‘minor’ (<5% PVE, 6 minor QTL). Interestingly, only one of the three glume YR QTL corresponded to a ‘major’ foliar YR locus: *QYrg.niab-2D.1* (glume 3.9% PVE) versus *QYr.niab-2D.1* (leaf 18% PVE). The finding that the remaining two glume resistance loci corresponded to minor foliar leaf QTL, and that none of the remaining seven ‘major’ foliar resistance loci (each controlling 6-17% foliar PVE) were relevant in the glumes, was somewhat unexpected. Together this indicates that while genetic mechanisms are shared between glume and leaf YR resistance, the importance of specific genetic loci in controlling these resistances is different between the two tissues. This finding has implications for breeding targets for current agricultural environments conductive to late *Pst* pressure, for environments likely to become higher risk due to changes in climate, or for the use of earlier flowering varieties.

Literature searches found all three of our glume/leaf resistance loci to have named *Yr* resistance genes on relevant chromosome arm, and alien chromosome introgression was implicated as underlying the source of resistance in two of the three loci. The physical map location of our glume/leaf QTL on the long arm of chromosome 5A overlapped with named adult plant resistance locus *Yr34*, which has recently been shown to have originated from *T. monococcum* ssp. *monococcum* (Chen et al. 2021). Predicted allelic effects at our chromosome 5A glume/leaf locus showed alleles from founders Alchemy, Brompton, Claire and Hereward conferred resistance. Our bioinformatic analysis of six published wheat genome assemblies included the variety Claire, which showed it to carry a putative alien introgression (at ~703.0-708.5 Mbp) within the QTL interval. The introgression is particularly common in European wheat, and cvs. Alchemy, Brompton and Claire were all recently reported to carry the introgression, from position 703.0 Mbp to the telomere (Chen et al. 2021; although it was not stated whether Hereward had been screened). Collectively, these data indicate *Yr34* may underlie our co-locating glume and foliar YR resistance locus *QYrg.niab-5A.1/QYr.niab-5A.1*, and that *Yr34* still acts as an effective quantitative source of YR resistance in modern wheat, despite its introgression into wheat more than 200 years ago (Chen et al. 2021). For our glume/leaf QTL *QYrg.niab-2D.2/QYr.niab-2D.1*, adult plant resistance gene *Yr54* (at which resistance is conferred by alleles from the Mexican wheat variety Quaiu 3; Banset et al. 2014) was predicted to lie within our QTL interval. Additionally, our 2D locus overlapped with the adult plant YR resistance QTL *QPst.jic-2D.1* (based on marker gwm301), for which resistance is conferred by the German variety Alcedo (Jagger et al. 2011). YR glume and leaf resistance in our MAGIC population was conferred at the chromosome 2D locus by alleles originating from Alchemy and Claire. Indeed, pedigree analysis shows Alchemy is a progeny line developed from a cross using Claire. No pedigree link has been reported between Quaiu 3 and Alcedo (Jagger et al. 2011), and while Alcedo is in the pedigree of some UK varieties (Fradgley et al. 2019), it has no clear link to Alchemy or Claire. Interestingly, our analysis of genomic sequences of six wheat varieties found evidence in the MAGIC founder Claire (and cv. Weebill1) of an alien introgression on chromosome 2D (from ~635.5-641.0 Mbp in the reference genome) that spans our 2D QTL peak (at 638.4 Mbp). As (i) alleles from Claire and Alchemy conferred glume/leaf resistance at the *QYrg.niab-2D.2/QYr.niab-2D.1* locus, and (ii) Claire is a parent of Alchemy, and (iii) Claire and Alchemy carry an identical haplotype across our 2D locus based on 90k SNP array data (Bouvet et al. 2021b), this indicates that resistance at this locus may conferred by alleles carried by an alien chromosomal introgression. Collectively, these results support the hypothesis that the sources of resistance for our chromosome 5A YR QTL is due to is due to *Yr34*, while that from our 2D QTL is from an unknown alien chromosomal introgression.

### A role for phenology in glume YR resistance

Ear infection by *Pst* commonly affects susceptible wheat varieties. However, YR ear infection of relatively resistant wheat varieties has been reported in New Zealand (Cromey, 1989b). This phenomenon was observed in the MAGIC founder Soissons, which was very susceptible to *Pst* glume infection (Figure 1c) but had very high leaf resistance (Bouvet et al. 2021b). While this difference could be due to several factors, Soissons is notable amongst the MAGIC founders for its early flowering phenotype - due largely to carrying the photoperiod insensitive early-flowering *Ppd-D1a* allele. The most significant SNP for the glume YR QTL *QYrg.niab-2D.1* was within the *Ppd-D1* gene, and represents a diagnostic marker that discriminates between the early flowering Soissons allele and the late-flowering allele(s) carried by the remaining seven MAGIC founders at this gene. As the early *Ppd-D1a* allele was associated with increased glume YR susceptibility, early flowering likely effects glume susceptibility pleiotropically. Further evidence of a link between YR ear infection and flowering time was provided by the co-location of *QYrg.niab-4A.1* with the multi-year, multilocation flowering time QTL *QFt.niab-4A.03*, previously identified in the same MAGIC population (Ian Mackay, personal communication). An influence of flowering time on glume YR infection was somewhat expected, as flowering time often influences other fungal diseases of the wheat ear. While earlier flowering was associated with increased glume YR infection, early varieties often come with other benefits. These include avoidance of fungal diseases of the ear such as fusarium head blight (Gervais et al. 2003; Goddard et al. 2021) and Septoria nodorum glume blotch (Aguilar et al. 2005; Downie et al. 2020; Lin et al. 2020a, 2020b; Scott et al. 1982), as well as drought avoidance. Early flowering can also be beneficial due to agronomic considerations, such as the timing and logistics of harvest.

Therefore, despite evidence UK wheat varieties released over the last 10 years have an increasingly narrower range of flowering time with a much smaller proportion of early varieties currently available (Sheenan & Bentley, 2020), there is still demand for early flowering varieties - which accounted for 14% of certified UK seed weight in 2019 (NIAB TAG, 2020).

### Conclusions

While we analysed just two trials for the identification of genetic loci controlling glume YR infection, this study nevertheless provides a starting point towards understanding the genetic components underlying YR resistance in wheat glumes, how it relates to foliar resistance, and demonstrates that subsets of genetic loci controlling flowering time likely also have a pleiotropic effect. The knowledge of which QTL are relevant to both leaf and glume resistance will help marker-assisted breeding approaches develop wheat varieties best suited for cultivation in agricultural environments in which glume *Pst* infection is prevalent.

## Abbreviations

BLUEs: Best Linear Unbiased Estimators
MAGIC: Multi-parent advanced generation inter-cross
MAS: Marker-assisted selection
*Pst*: *Puccinia striiformis* Westend f. sp. *tritici*
QTL: Quantitative trait locus
RIL: Recombinant inbred line
SNP: Single nucleotide polymorphism
YR: Yellow rust

## Acknowledgements

LB was funded via a Biotechnology and Biological Sciences Research Council (BBSRC) Doctoral Training Partnership PhD award to the University of Cambridge and undertaken at NIAB. LB also received funding from the Roger Harrison Trust towards travel expenses. JC’s time was supported in part by Biotechnology and Biological Sciences Research Council (BBSRC) grant BB/P010741/1. We thank Prof Julian Hibberd (University of Cambridge) for supervisory inputs within the Doctoral Training Programme, Dr Keith Gardner (NIAB) for genetics training, and the NIAB Trials Team for the sowing, growth and agronomic care of the NIAB field trail.

## Author contributions

LB carried out trial preparation, pathology work, yellow rust assessments and all statistical and genetic analyses. SB and PW managed the trial at Osgodby. LP undertook genomic analyses. IM, SH and JC provided project supervision and resources. LB and JC wrote the manuscript. All authors edited and approved the manuscript.

## Notes

### Competing Interest Statement

The authors have declared no competing interest.

