## Supplementary Figure 1 for "Genetic resistance to stripe rust infection of the wheat ear is controlled by genes controlling foliar resistance and flowering time"

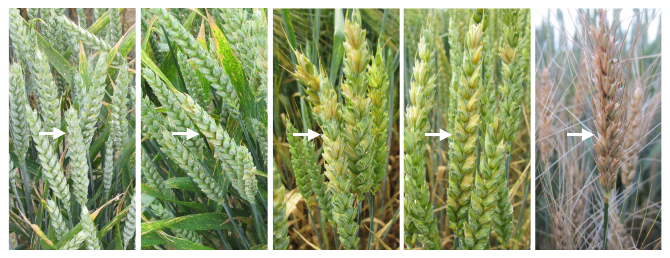

**Supplementary Figure 1.** Examples of the glume yellow rust infection scoring system used. From left to right: score 0 (no glume infection), score 1 (25% of glumes infected), score 2 (50% of glumes infected), score 3 (75% of glumes infected), score 4 (all glumes per head infected).
