## Supplementary Figure 2 for "Genetic resistance to stripe rust infection of the wheat ear is controlled by genes controlling foliar resistance and flowering time"

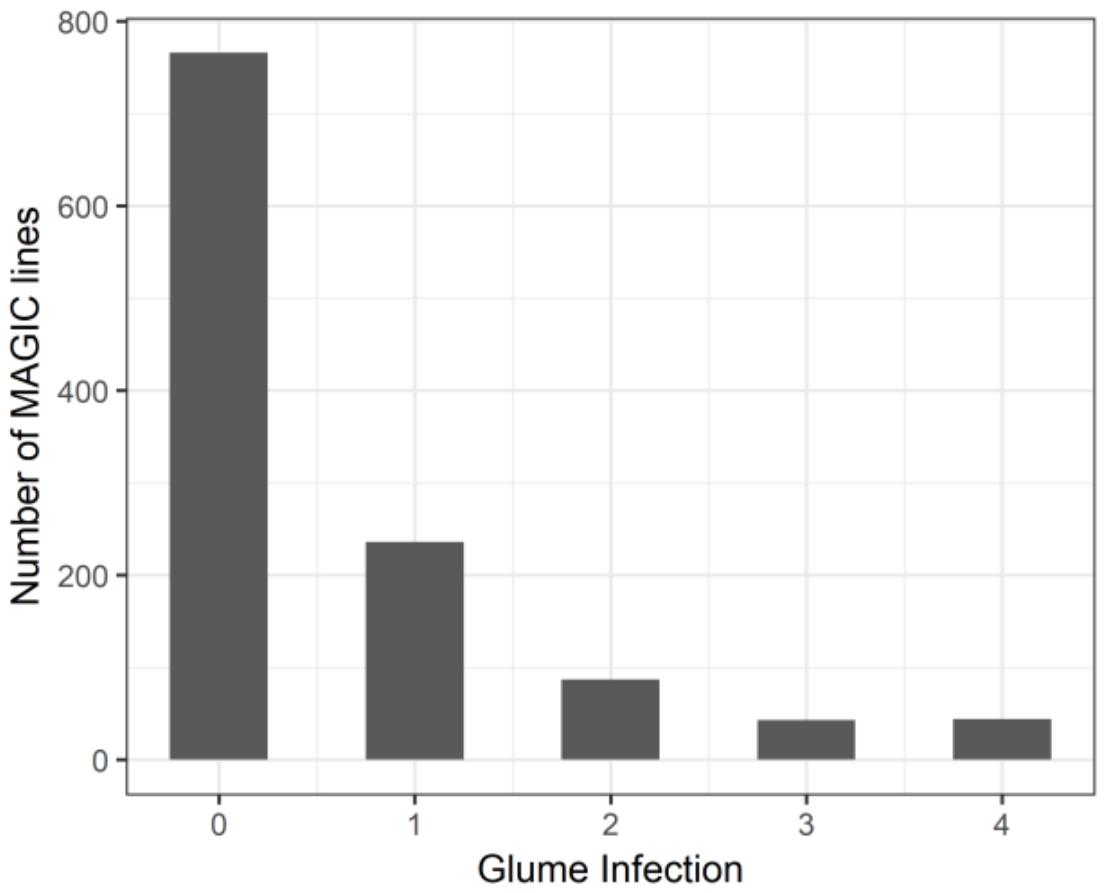

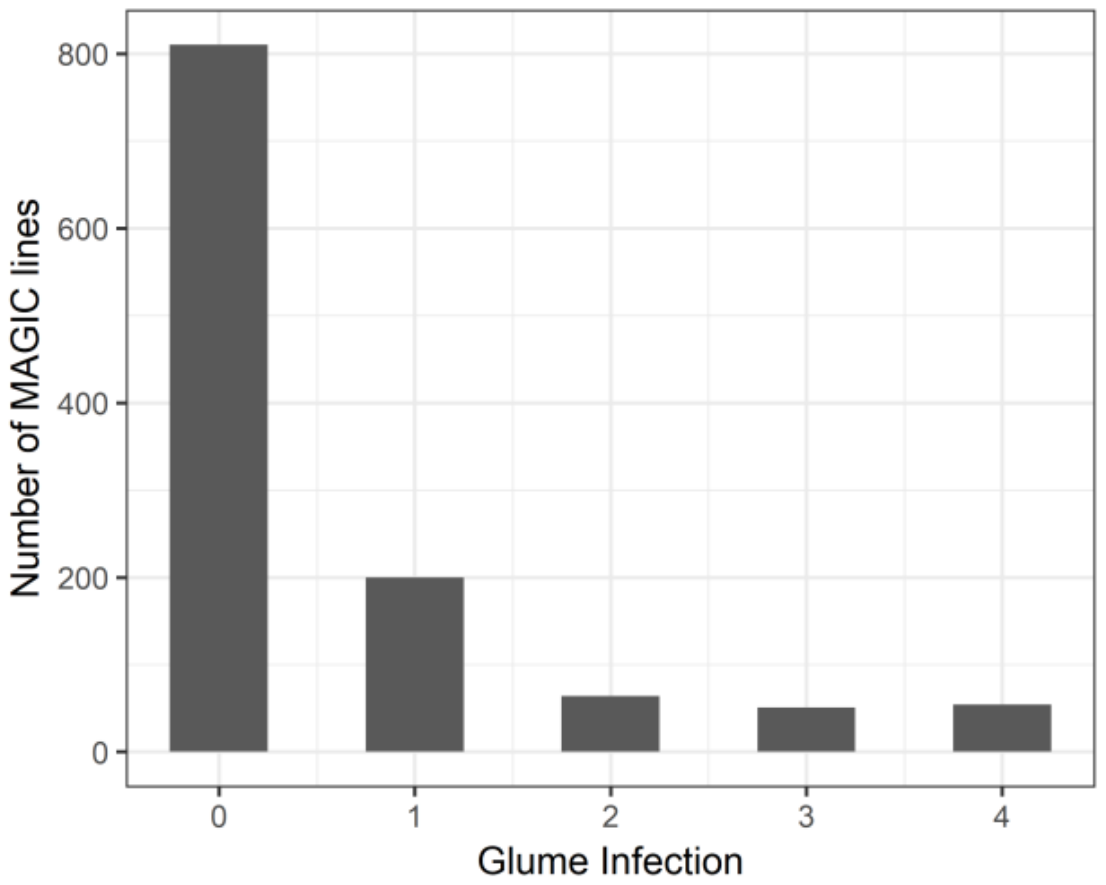

**Supplementary Figure 2.** Histogram of yellow rust glume infection in the MAGIC recombinant inbred lines (RILs) at trials NIAB16 (left) and OSG16 (right). The number of MAGIC RILs includes single counts for un-replicated lines and total counts for replicated lines.
