## Supplementary Figure 3 for "Genetic resistance to stripe rust infection of the wheat ear is controlled by genes controlling foliar resistance and flowering time"

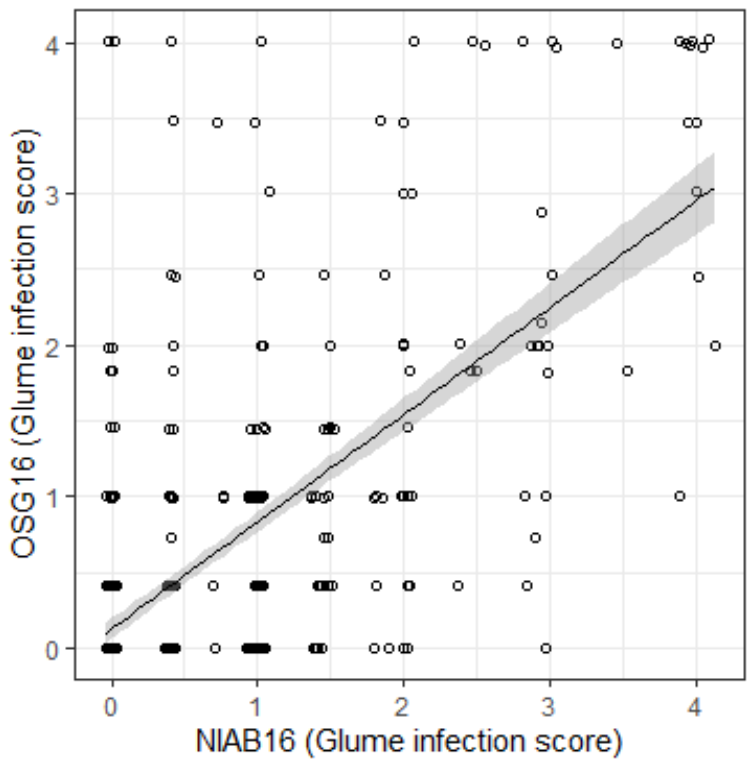

**Supplementary Figure 3.** Scatter plot showing the correlation between glume yellow rust infection at trial sites NIAB16 and OSG16. Scatter plot shows back-transformed adjusted means. The black line is the linear regression line and the grey area the 95% confidence level interval. Correlation coefficients: *R* = 0.47, *p* <2.2E-16.
