## Supplementary Figure 4 for "Genetic resistance to stripe rust infection of the wheat ear is controlled by genes controlling foliar resistance and flowering time"

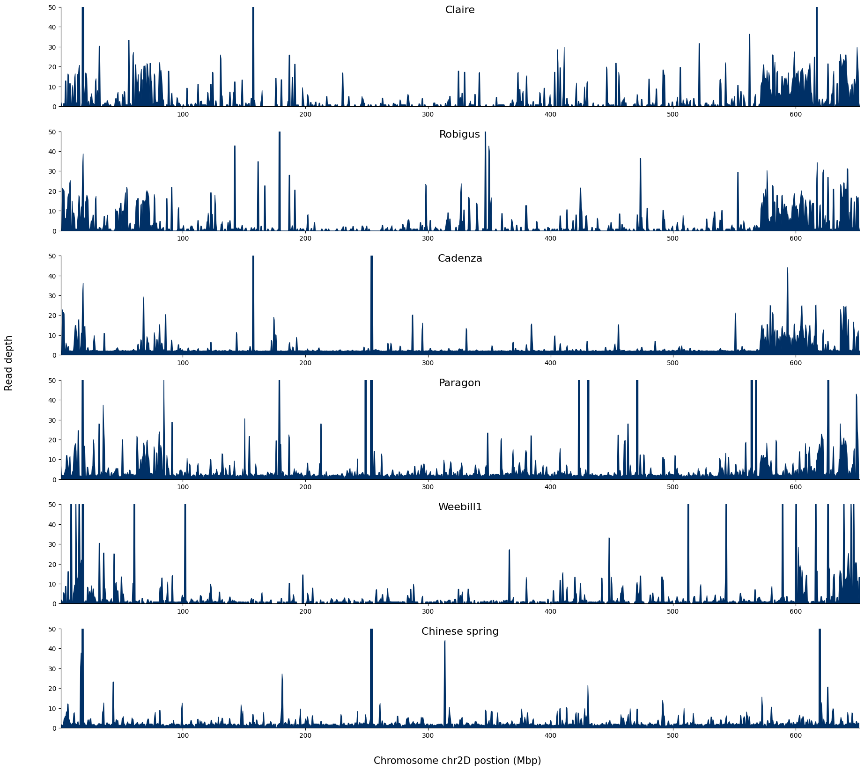
a

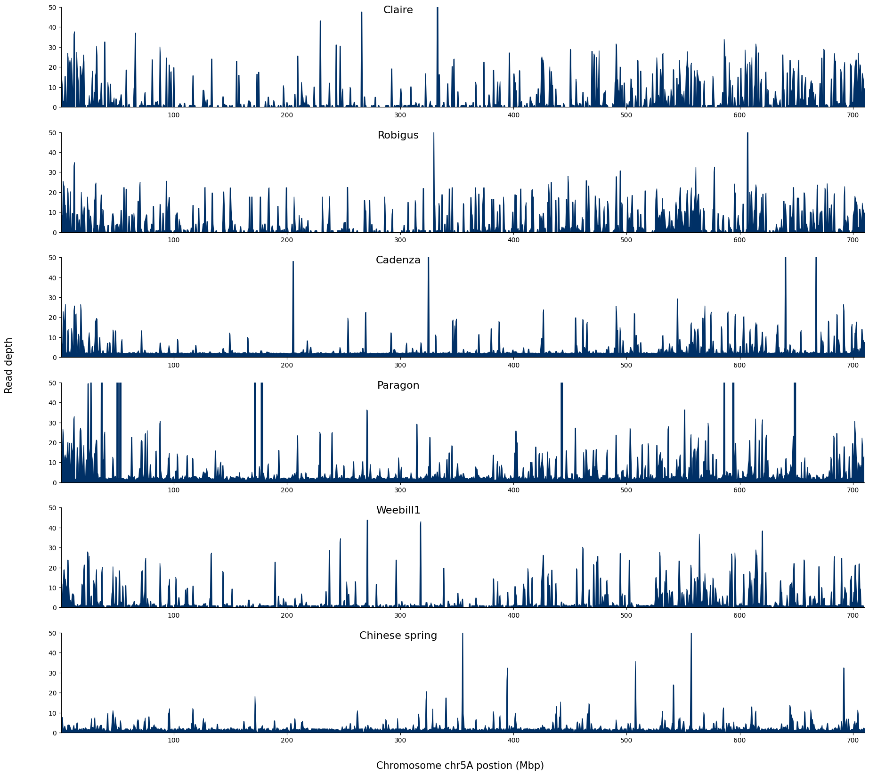

b

**Supplementary Figure 4.** Alignment depths from the self-alignment of wheat cultivars reads unmapped in the Chinese Spring reference, anchored back to chromosomes 2D and 5A of the Chinese spring reference genome using the gene to scaffold alignment information generated in our minimap2 based diversity analysis. Plots are capped to a read depth of 50. An increased depth can be seen at regions previously identified as being of low depth in BWA mem alignments and higher diversity in our diversity analysis.
