## Supplementary Table 1 for "Genetic resistance to stripe rust infection of the wheat ear is controlled by genes controlling foliar resistance and flowering time"

| **Location** | **Isolate code** | **Year** | **Pathotype** | **Virulence profile** |
| --- | --- | --- | --- | --- |
| Cambridgeshire | 16/009 | 2016 | Red race 5 (Warrior group) | 1,2,3,4,6,7,9,17,25,32,Re,Sp,Ro,So,Ca,Ap |
| Cambridgeshire | 16/048 | 2016 | Pink (Path) race 13 (Warrior group) | 1,2,3,4,6,7,9,17,25,32,Re,Sp,Ro,So,Wa,Ca,(Ap) |
| Cambridgeshire | 16/199 | 2016 | Red race 5 (Warrior group) | 1,2,3,4,6,7,9,17,25,32,Re,Sp,Ro,So,Ca,Ap |
| Cambridgeshire | 16/204 | 2016 | Red race 11 (Warrior group) | 1,2,3,4,6,7,9,17,25,32,Re,Sp,Ro,So,(Wa),Ca,St,Ap |
| Cambridgeshire | 16/205 | 2016 | Red race 5 (Warrior group) | 1,2,3,4,6,7,9,17,25,32,Re,Sp,Ro,So,Ca,Ap |
| Cambridgeshire | 16/208 | 2016 | Blue race 7 (Warrior group) | 1,2,3,4,6,7,9,17,25,32,Re,Ro,So,Ev |
| Lincolnshire | 16/131 | 2016 | Red race 24 (Warrior group) | 1,2,3,4,6,7,9,17,25,32,Re,Sp,Ro,So,Wa,Ca,St,Ap |
| Lincolnshire | 16/286 | 2016 | Red race 23 (Warrior group) | 1,2,3,4,6,7,9,17,25,32,Re,Sp,Ro,So,Wa,Ca,Ap |
| Lincolnshire | 16/288 | 2016 | Red (Path) race 5 (Warrior group) | 1,2,3,4,6,7,9,17,25,32,Re,Sp,Ro,So,Ca,(Kr),Ap |
| Lincolnshire | 16/289 | 2016 | Red race mix (Warrior group) | 1,2,3,4,6,7,8,9,17,25,32,Re,Sp,Ro,So,Wa,Ca,St,Kr,Ap |
| Lincolnshire | 16/290 | 2016 | Red race 23 (Warrior group) | 1,2,3,4,6,7,9,17,25,32,Re,Sp,Ro,So,Wa,Ca,Ap |
| Lincolnshire | 16/292 | 2016 | Pink race 14 (Warrior group) | 1,2,3,4,6,7,9,17,25,32,Re,Sp,Ro,So,Wa,Ca,St,Kr,Cr |
| Osgodby, Lincolnshire | 16/277 | 2016 | Blue race 10 (Warrior group) | 1,2,3,4,6,9,17,25,32,Re,Sp,Ro,So,Wa |

**Supplementary Table 1.** Summary of *Pst* pathotypes identified from samples collected and tested by the UK Pathogen Virulence survey from Cambridgeshire and Lincolnshire in 2016 (Hubbard et al. 2017). Virulence profile corresponds to virulence on yellow rust resistance genes *Yr1, Yr2, Yr3, Yr4, Yr6, Yr7, Yr8, Yr9, Yr17, Yr25, Yr27, Yr32* and on the varieties Rendezvous (Re), Spaldings Prolific (Sp), Robigus (Ro), Solstice (So), Warrior (Wa), Ambition (Am), Cadenza (Ca), KWS Sterling (St), Kranich (Kr), Apache (Ap), Crusoe (Cr), Evolution (Ev) and Timber (Ti). Brackets indicate a borderline virulent response.
