## Supplementary Table 3 for "Genetic resistance to stripe rust infection of the wheat ear is controlled by genes controlling foliar resistance and flowering time"

| Site | REML estimates | | | | | | | σ^2^e | *h*^2^ |
| --- | --- | --- | --- | --- | --- | --- | --- | --- | --- |
|  | Random |  |  |  | Fixed |  |  |  |  |
|  | Term | Est. | SE |  | Term | Wald | *Chi pr* |  |  |
| NIAB16 | column | 0.00137 | 0.0013 |  | G | 4227.1 | <0.001 | 0.0603 | 0.80 |
| OSG16 | rep.block | 0.00009 | 0.00043 |  | G | 4773.1 | <0.001 | 0.0568 | 0.82 |

**Supplementary Table 3.** Summary of Restricted Maximum Likelihood (REML) covariance estimates of the mixed linear model for glume yellow rust resistance in the MAGIC population at trial sites NIAB16 and OSG16. Est.: estimates; SE: standard error; G: genotype; *h*^2^: broad sense heritability; σ^2^e: residual error.
